# Structural Insights into Native Intact *Mycobacterium abscessus* by Conventional and Ultrahigh-field solid-state NMR at 1.2 GHz

**DOI:** 10.64898/2026.05.19.726312

**Authors:** Chang-Hyeock Byeon, Yu-Hao Wang, Abdulkadir Tunc, W. Trent Franks, William H. DePas, Ümit Akbey

## Abstract

We present an ultrahigh-field magic-angle spinning (MAS) solid-state NMR (ssNMR) study to characterize intact nontuberculous mycobacteria (NTM) at the molecular level. Hydrated and dried whole-cell *Mycobacterium abscessus* samples were investigated by combining conventional high-field ssNMR at 750 MHz with ultrahigh-field ssNMR at 1.2 GHz and ultrafast MAS at 100 kHz. To improve sensitivity and enable multidimensional experiments, 13C/15N isotope labeling was performed after growth in synthetic cystic fibrosis medium (SCFM). We utilized 1D 13C and multidimensional 1H-13C and 13C-13C ssNMR experiments to characterize the chemical composition, dynamics, and structural organization of the *M. abscessus* cell envelope. The isotope-labeling efficiency was found to be non-uniform across different molecular classes, with high incorporation into polysaccharides and lower incorporation into lipid and peptide-associated signals. INEPT- and CP-based experiments selectively probed flexible and rigid fractions of the samples, revealing substantial differences in linewidth, dynamics, and sensitivity between hydrated and dried preparations. Conventional 750 MHz experiments provided high-resolution multidimensional spectra and enabled identification of distinct chemical environments associated with peptidoglycan, arabinogalactan, mycolic acids, lipids, and peptide-associated components. Ultrahigh-field ssNMR at 1.2 GHz combined with ultrafast MAS and 1H detection substantially improved spectral resolution and sensitivity in particular per mg of sample amount, allowing detection of weak and previously unresolved resonances, including polysaccharide and possible nucleic-acid-associated signals. Together, these results demonstrate that ultra-high-field and ultrafast-MAS ssNMR enables detailed characterization of intact NTM cell envelopes under near-native conditions and provides a framework for future molecular investigations of antimicrobial interactions.

## 1. Introduction

Antimicrobial resistance (AMR) is increasingly compromising the effectiveness of established therapies, particularly in chronic infections that require prolonged treatment and thereby accelerate the emergence of resistant strains.^1^ Within this context, NTM have emerged as clinically significant opportunistic pathogens characterized by intrinsic tolerance to many conventional antibiotics and steadily increasing prevalence.^2-4^ In the United States, infections caused by NTM have surpassed those associated with *Mycobacterium tuberculosis* for more than a decade,^3^ and their incidence continues to rise.^4^ These organisms predominantly cause pulmonary disease, especially in individuals with underlying lung disorders such as cystic fibrosis (CF) and non-CF bronchiectasis.^5-7^ Among the NTM able to cause disease in humans, *Mycobacterium abscessus* represents a major clinical challenge due to its pronounced drug resistance and poor treatment outcomes, with failure rates approaching 75% in some patient cohorts. ^8-13^

A central determinant of this behavior is the unique structure of the mycobacterial cell envelope. In contrast to classical Gram-positive and Gram-negative bacteria, NTM possess a multilayered envelope composed of peptidoglycan (PG) covalently linked to arabinogalactan (AG) and coated by a lipid-rich outer membrane known as the mycomembrane (MM).^14-16^ This outer layer, dominated by long-chain mycolic acids (MA) and supplemented by diverse lipids, polysaccharides, and proteins, forms a dense and highly impermeable hydrophobic barrier. The chemical composition, spatial organization, and dynamics of this envelope govern key biological processes, including antibiotic tolerance, host-pathogen interactions, and biofilm formation.^17-21^ Despite its importance, a detailed molecular-level understanding of the intact envelope remains limited, largely due to its chemical complexity, heterogeneity, and the coexistence of rigid and highly dynamic components.

Solid-state NMR (ssNMR) spectroscopy provides a uniquely suited approach for probing such complex biological systems. Unlike solution-based techniques that rely on solubilization or chemical extraction, ssNMR enables direct and quantitative characterization of intact, heterogeneous materials under near-native conditions.^22-27^ This capability is particularly important for systems such as whole cells and biofilms, where both soluble and insoluble components contribute to structure and function. In addition, ssNMR can selectively probe molecular dynamics through different polarization transfer mechanisms, allowing differentiation between rigid (via CP ssNMR) and flexible (via INEPT ssNMR) fractions within the same sample.^28-34^ However, the application of ssNMR to whole-cell systems is often limited by spectral overlap, restricted chemical shift dispersion at conventional magnetic field applications, and sensitivity constraints especially at natural isotopic abundance such as in samples extracted from patients. While multidimensional (nD) ssNMR experiments can partially alleviate these challenges, they frequently require long acquisition times or isotopic labeling strategies that are not always feasible for native or clinically relevant samples.^28-39^ Nevertheless, for lab-grown NTM, isotope-labelling is feasible and provide the possibility of recording nD ssNMR spectra for better resolution,^40^ which is still a promising path in addition to the natural-abundance native (e.g. patient) NTM sample characterization.

Recent advances in NMR instrumentation offer a pathway to overcome these limitations. In particular, the combination of ultrahigh magnetic fields above 1 GHz and ultrafast magic-angle spinning (MAS) above 100 kHz significantly enhances both spectral resolution and sensitivity. Higher magnetic fields increase chemical shift dispersion and reduce overlap between resonances, while ultrafast MAS efficiently averages residual dipolar interactions, leading to narrower linewidths, especially in proton-detected (^1^H-detected) experiments. Together, these developments enable access to previously unresolved spectral features in complex biological systems and facilitate multidimensional experiments within practical timeframes. Combined with the isotope-labelling possibility for lab-grown NTM, this experimental setup is highly informative and could also facilitate more difficult spectral analysis of patient-extracted samples.

In our previous work, we demonstrated that hydrated whole-cell NTM can be investigated by ssNMR under near-native conditions without isotope-labelling,^41^ enabling identification of major chemical components and providing insight into their structural organization and dynamics. Along with the few other ssNMR investigations on NTM,^40, 42^ this work constitutes a basis for structural characterization by ssNMR. These studies established the feasibility of whole-cell ssNMR for NTM and revealed distinct contributions from lipids, polysaccharides, and protein-associated components. At the same time, they highlighted limitations arising from spectral overlap reducing resolution under conventional field strengths and MAS conditions, which restrict the level of molecular detail that can be extracted.

Here, we build on these previous work and expand it by applying ultrahigh-field ssNMR at 1.2 GHz in combination with ultrafast MAS at 100 kHz to intact native whole-cell NTM samples that are uniformly 13C,15N isotope-labeled. By performing experiments at ultrahigh magnetic field and ultrafast MAS frequency, we achieve improvements in both resolution and sensitivity, enabling separation of overlapping resonances and allow more detailed characterization of cell-envelope components. This is in particular true for the dry sample suffering the most from line-broadening, which was utilized as a means of sensitivity increase (>10x compared to the hydrated wet sample). Moreover, sample drying also provides means to detect mobile chemical species in addition to the rigid ones at once more quantitatively, compared to hydrated sample where the flexible and rigid fractions appear in different ssNMR spectra (INEPT vs. CP). This approach allows us to probe molecular composition and dynamics of planktonic *M. abscessus* with high sensitivity and resolution that are not accessible under conventional conditions, in particular when the sample amount is limited. Overall, our results demonstrate that ultrahigh-field ssNMR, ultrafast MAS and 1H-detection combined with isotope-labelling and conventional nD ssNMR, could transform the study of complex bacterial systems and provides a powerful platform for resolving the molecular architecture underlying antimicrobial tolerance in NTM.

## 2. Methods

### Bacterial strains and sample preparation

*Mycobacterium abscessus* 0253a, a smooth colony isolate from a person with CF,^43^ was grown in TYEM plus 0.05% Tween-80 for 48-72 hours at 37°C, with shaking at 250 RPM.^44^ For preparation of samples for ssNMR analysis, 0253a stationary phase cultures were diluted into 250 mL of SCFM to a final optical density (OD_600_) of 0.01. The composition of SCFM was the same as described in Palmer et al. with some modifications.^45^ The final concentration of Morpholinepropanesulfonic acid (MOPS) was increased from 10mM to 20mM in order to increase the buffer capacity. The base SCFM contains 3mM D-glucose and 2.3mM NH_4_+. Here, SCFM contained 13mM of isotope-labeled D-Glucose-^13^C_6_ (Sigma-Aldrich, 660663) and 2.3mM of isotope-labeled ammonium-^15^N chloride (Sigma-Aldrich, 299251). This recipe allows for detergent-free planktonic growth of *M. abscessus*.^43^ The 250 mL cultures were grown at 37°C with shaking at 250 RPM. The cells were harvested at 60 hours post-inoculation by concentrating the cell culture in a centrifuge at 7,000 x g for 10 minutes at 4°C. The resulting pellet was resuspended in 30 mL 1x PBS with 4% paraformaldehyde (PFA) and fixed overnight at 4°C. The PFA-fixed cells were then centrifuged again at 7,000 x g for 10 minutes at 4°C. The resulting pellet was resuspended in 1.5 mL ice-cold 1x PBS and transferred to a new Eppendorf tube and prepped for ssNMR analysis.

### ssNMR spectroscopy

ssNMR experiments were performed at two magnetic field strengths. Conventional-field measurements were carried out at a 1H Larmor frequency of 750 MHz on a Bruker Avance III spectrometer equipped with a low-temperature triple-resonance 3.2 mm MAS probe. Thin-walled 3.2 mm rotors were used, with ∼40-50 mg of sample. Ultrahigh-field experiments were conducted at a 1H Larmor frequency of 1.2 GHz on a Bruker Avance NEO spectrometer equipped with a 0.7 mm triple-resonance ultrafast MAS probe, with ∼0.5 mg of sample. The 3.2 mm MAS probe at 1.2 GHz was not utilized and compared to the results.

Hydrated intact *M. abscessus* whole-cell planktonic samples were packed into MAS rotors by transferring hydrated pellets directly into the rotor body using pipette tips, followed by centrifugation using a benchtop centrifuge. For all MAS experiments, temperatures were actively regulated to maintain near-ambient sample conditions and to ensure physiologically relevant states throughout data acquisition. ^13^C shifts were externally referenced by adjusting the adamantane CH_2_ peak to 38.48 ppm following the TMS scale.^46^ The chemical shift referencing were confirmed by using the 0 ppm 1H DSS reference value that is added to the samples, and by referencing 13C chemical shifts indirectly via the 1H frequency.^47^

### ssNMR experiments at 750 MHz and 10 kHz MAS

1D / 2D ssNMR experiments at 750 MHz were performed as follows. 1D 13C CP/INEPT and 2D 1H-13C CP and CP/INEPT spectra were acquired at an MAS frequency of 10 kHz MAS. π/2 RF pulse lengths for 1H: ∼3.3 μs, 13C: ∼5 μs and 15N: ∼6 μs. CP experiments employed a 1 ms contact time with a 70–100% amplitude ramp on the 1H channel. Proton decoupling at approximately 90 kHz was applied during signal acquisition. 1D 13C spectra were acquired using INEPT, CP, and direct polarization (DP) experiments in same total acquisition times per dataset. 1024 scans with 1s recycle delays were used for CP and INEPT experiments, while a 204 scans with 5 s recycle delay were used for DP. 1D spectra were processed using Gaussian apodization with a broadening of 35 Hz (GB = 0.055) in TopSpin. The 1D NCa and NCO experiments were recorded following to standard conditions.^48^

1H-detected 2D 1H-13C INEPT/CP spectra on wet sample were acquired at 10 kHz MAS using 1024/512 transients, respectively, with 64 indirect-dimension increments and 1 s recycle delay (total acquisition times of ∼18/9 h for INEPT/CP). 13C-detected 2D 1H-13C INEPT/CP spectra on wet sample were acquired at 10 kHz MAS using 32/16 transients, respectively, with 64 indirect-dimension increments and 1 s recycle delay (total acquisition times of ∼0.5/0.25 h for INEPT/CP). 13C-detected 2D 1H-13C CP spectra on dry sample were acquired at 10 kHz MAS using 16 transients, respectively, with 64 indirect-dimension increments and 1 s recycle delay (total acquisition times of ∼0.25 h for INEPT/CP). 2D spectra were processed using Gaussian apodization (35 Hz, GB = 0.055) in the direct dimension and a mixed sine-squared window function (SSB = 3) in the indirect dimension. The contours of 2D spectra were consistently set to levels where the noise starts to appear.

### ssNMR experiments at 1.2 GHz and 100 kHz MAS

Ultrahigh-field ssNMR experiments were performed at 1.2 GHz and under ultrafast MAS at frequencies up to 100 kHz. 1H-detected experiments were used exclusively to capitalize on the enhanced sensitivity and resolution afforded by the combination of ultra-high magnetic field strength and ultrafast MAS. π/2 RF pulse lengths were calibrated on each sample with 1H: ∼1.6 μs, 13C: ∼3 μs and 15N: ∼3 μs, respectively. Low-power WALTZ16 heteronuclear decoupling schemes were employed during acquisition at 12.5 kHz. The nominal sample temperatures were ∼268 K and controlled.

2D 1H-13C correlation spectra were acquired using INEPT- and CP-based pulse sequences optimized for ultrafast MAS conditions. A recycle delay of 1.5 s was used, and the number of transients and indirect-dimension increments was adjusted to balance sensitivity, resolution, and experimental duration. 1H-detected 2D 1H-13C INEPT/CP spectra on wet sample were acquired at 100 kHz MAS using 4/64 transients, respectively, with 256 indirect-dimension increments (total acquisition times of ∼0.4/7 h for INEPT/CP). 1H-detected 2D 1H-13C CP spectra on dry sample were acquired at 100 kHz MAS using 32 transients, respectively, with 256 indirect-dimension increments (total acquisition times of ∼3.5 h for INEPT/CP). 2D spectra were processed using Gaussian apodization (35 Hz, GB = 0.015) in the direct dimension and a mixed sine-squared window function (SSB = 2) in the indirect dimension. The enhanced chemical-shift dispersion at 1.2 GHz, together with efficient averaging of anisotropic interactions at 100 kHz MAS, resulted in a reduction of spectral overlap compared to measurements performed at 750 MHz and slower MAS frequencies. Chemical shifts were externally referenced using standard protocols mentioned above. The contours of 2D spectra were consistently set to levels where the noise starts to appear.

## 3. Results

### 3.1. Structural organization of NTM and prior ssNMR characterization

The functional properties of NTM are closely linked to the molecular architecture of the cell envelope, which differs from that of classical bacterial systems, **Fig. 1**. This envelope consists of a PG scaffold covalently linked to AG and overlaid by a lipid-rich outer layer dominated by long-chain MA. This organization forms a highly effective hydrophobic permeability barrier that regulates environmental interactions and contributes to antimicrobial resistance.^14, 17^-^21, 49^ At the molecular level, the envelope is both chemically diverse and dynamically heterogeneous, comprising rigid structural elements alongside more flexible components

This chemical complexity is directly reflected in ssNMR spectra, where distinct chemical shift regions correspond to the major molecular classes, **Fig. 1**,**2**. Carbonyl signals span ∼190-160 ppm, aromatic and nucleic acid (NA, if present) contributions appear between ∼160-110 ppm, and polysaccharides, including PG and AG, dominate the ∼110-50 ppm region.^40-42^ Aliphatic signals from lipids and MA are observed between ∼50-10 ppm, with characteristic sharp methylene and olefinic resonances near ∼30 ppm and ∼130 ppm, respectively. The coexistence of these chemically and dynamically distinct components leads to substantial spectral overlap, representing a major limitation for identifying and quantifying individual molecular contributions, in particular with 1D ssNMR methods.

**Figure 1.**
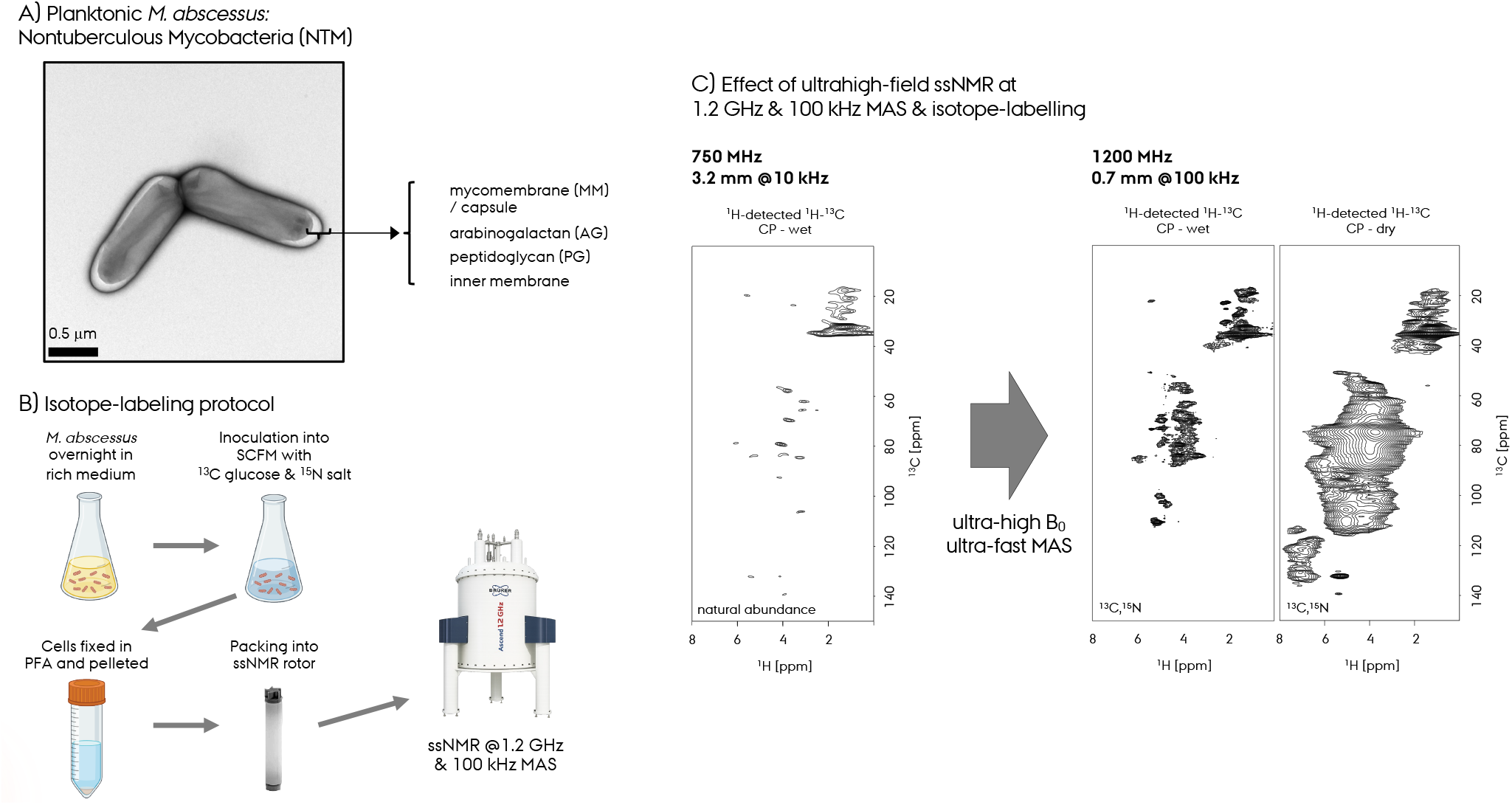
**A.** Negative-staining EM micrograph of *M. abscessus* along with a cartoon representation of the cell envelope architecture and its components: inner membrane, peptidoglycan (PG), arabinogalactan (AG), and mycomembrane (MM)/capsule. **B**. Protocol for ^13^C,^15^N isotope-labelling of *M. abscessus*. **C**. Ultrahigh-field ssNMR spectroscopy on wet and dry isotope-labeled *M. abscessus* at conventional 750 MHz (for natural abundance sample) and ultrahigh-field 1.2 GHz and 100 kHz (for ^13^C,^15^N sample).

Whole-cell and chemically extracted cell-envelope preparations each provide complementary advantages for ssNMR analysis.^40-42^ Intact whole-cell samples preserve native molecular organization and hydration, enabling characterization of physiologically relevant dynamics and intermolecular interactions. In contrast, extracted fractions simplify spectral interpretation and facilitate assignment of major envelope constituents, although chemical extraction can perturb native structural organization and selectively enrich or deplete specific components.

Pioneering ssNMR study by Cegelski and coworkers established a direct CP-based 1D ssNMR approach to probe natural-abundance mycobacterial cell-wall composition, for mAGP (cell-envelope: MA + AG + PG) and whole-cell samples.^42^ This enabled differentiation of key components such as PG, AG, and MA, and revealing species-dependent quantitative differences that maybe linked to antibiotic tolerance. Additionally, a high-resolution study from Schanda and coworkers demonstrated that isotope-labelling, cryogenic MAS probe and optimized experimental schemes significantly enhance sensitivity.^40^ The utilization of high-resolution 1H-detected ssNMR measurements on purified cell-wall and intact bacterial systems facilitated nD experiments and resonance assignment. The latter study also emphasized the importance of combining polarization transfer approaches to selectively probe rigid and mobile fractions within structurally heterogeneous cell envelopes. As a follow up on these, we recently presented 1D and 2D 1H-13C ssNMR via 13C-detection of two hydrated whole-cell NTM, *Mycobacterium smegmatis* and *M. abscessus*, under native conditions without isotope labeling.^41^ We identified over ∼100 distinct carbon signals and determined numerous polysaccharide species. Our work further demonstrated that differences in flexibility of different chemical sites can be resolved, revealing that *M. abscessus* exhibits a more flexible and reduced peptidoglycan layer compared to *M. smegmatis*. Collectively, these establish ssNMR as a powerful platform for identifying cell-envelope composition and dynamics in NTM.

Despite these advances, spectral overlap and limited chemical-shift dispersion under conventional high magnetic fields of 600-950 MHz and sensitivity-optimized conditions (CryoMAS ssNMR probes and 1H-detection at ultrafast-MAS) continue to restrict detailed molecular interpretation of intact NTM systems. While ultra-high magnetic fields above 1 GHz provide measurable improvements in spectral resolution, manifested as linewidth narrowing and improved sensitivity, CryoMAS approaches primarily enhance sensitivity and do not fully resolve spectral overlap alone in heterogeneous samples. Here, we address these and compare conventional high-field ssNMR (750MHz with 3.2mm MAS rotors) and ultrahigh magnetic fields (∼1.2 GHz) at ultrafast MAS (>100 kHz). Altogether, combining these with isotope-labeling protocol, we now enable simultaneous gains in spectral resolution and sensitivity to access molecular complexity of intact whole-cell NTM cell envelopes under native conditions.

### 3.2. Isotope labeling efficiency on *M. abscessus* and conventional ssNMR

To improve sensitivity and enable nD experiments, 13C,15N isotope labeling of planktonic *M. abscessus* was performed under SFCM growth conditions. The whole-cell *M. abscessus* bacteria samples were PFA fixed before the ssNMR measurements, for the sake of NMR spectroscopists’ health. We showed that the fixation doesn’t affect mycomembrane structure much.^41^ This allows the recording of nD ssNMR spectra, including 13C-detected 2D 13C-13C spectra with increased efficiency, that is not feasible to record at natural abundance without DNP.^50^ First, conventional ssNMR at moderate MAS frequencies of 10 kHz was applied by packing the fixed post-harvested cells in PBS buffer by gentle centrifugation as wet and fully hydrated pellets without any further treatment. We recorded ssNMR spectra at close to native conditions, as well as dried samples to increase sensitivity. Compared to our previous natural-abundance NTM study, the sensitivity of these samples is superior.^41^

The incorporation of 13C and 15N labels was quantified by recording conventional ssNMR at 750 MHz, 3.2mm probe and 10 kHz MAS for natural abundance and isotope labeled samples and was found to be non-uniform across different chemical species **Fig. 2A**. Polysaccharide signals (PG and AG) exhibited high labeling, whereas peptide and lipid resonances showed low efficiency, reflecting differences in metabolic pathways and glucose precursor utilization, **Fig 2A,B**. The normalized isotope-labeled and unlabeled 13C CPMAS ssNMR spectra of the dried planktonic *M. abscessus* sample showed that the13C-labelling efficiency is ∼100% for PS and∼25% for lipids, ∼20% for CO, and 20% for the peptide Ca/Calip signals. The 13C INEPT ssNMR spectra of the same dried sample contains only the highly-dynamic lipid chemical groups with ∼25% isotope-labelling efficiency, supporting the efficiency in the CP experiments. Without isotope-labelling, the high-sensitivity 1D 13C ssNMR spectra were able to be acquired from fixed whole-cell samples (as it is hydrated/wet and dried) using approximately ∼40-50 mg of material within hours. With isotope-labelling, despite lower efficiency for certain chemical groups, the similar sensitivity 1D 13C ssNMR spectra could be obtained in minutes, allowing high-efficiency 2D/3D ssNMR spectroscopy. These isotope-labeling efficiency ratios will be crucial in determining the abundance of different chemical species from the ssNMR spectra, discussed below. The 15N isotope-labelling efficiency was lower and not determined due to sensitivity limitations.

**Figure 2.**
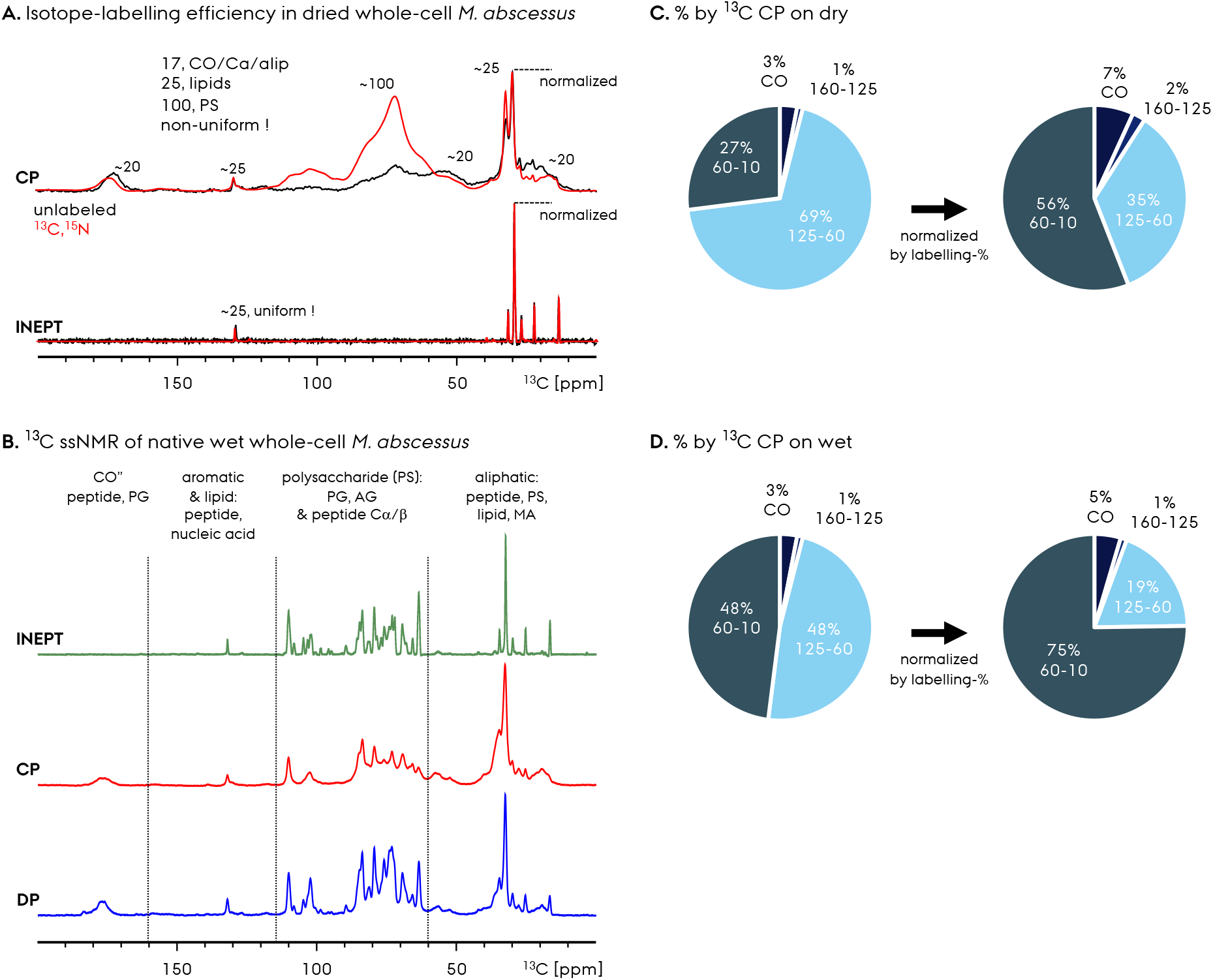
ssNMR spectra recorded on wet/dried unlabeled and ^13^C,^15^N isotope-labeled *M. abscessus* whole-cell samples at 750 MHz recorded with 3.2 mm thin-wall rotor and 10 kHz MAS. **A.** Comparison of 1D ^13^C CPMAS ssNMR spectra of dried unlabeled (black) and 13C,15N isotope-labeled (red) *M. abscessus* whole-cell samples. The spectra were recorded with CP and INEPT polarization transfer. **B**. Comparison of 1D ^13^C ssNMR spectra of wet ^13^C,^15^N isotope-labeled *M. abscessus* whole-cell samples with INEPT, CP and DP. Tentative chemical group assignments are shown. Quantification of different chemical shift ranges determined by using ^13^C spectra on **C**. dry and **D**. wet samples shown in A,B. The left plot shows the abundances determined directly from the CP spectra, whereas the right plot shows the normalized abundances calculated by using the isotope-labeling efficiencies that is ∼4-fold larger for the AGP fraction.

To capture the full range of molecular dynamics, complementary polarization schemes were employed, on wet hydrated native samples. INEPT-based 13C ssNMR experiments selectively detect flexible chemical components, while CP detects rigid regions of the cell envelope. In addition, DP experiments provide a combined view of both fractions and offer quantification possibility. Together, these approaches enable a complete characterization of chemical composition and dynamics, **Fig. 2B,C**. A pronounced difference in spectral resolution was observed between the flexible and rigid fractions obtained from the hydrated native whole-cell sample, **Fig. 2B**. These three types of 13C ssNMR spectra were recorded within the same total experiment time at 750 MHz, so the signal intensities are directly comparison. INEPT-based spectra (green) exhibited narrow linewidths, with average values of ∼70 Hz (∼0.4 ppm), whereas CP-based spectra (red) were substantially broader, with linewidths on the order of ∼200 Hz (∼1.1 ppm). These differences reflect the underlying molecular motion, with flexible species producing sharper resonances and rigid components giving rise to broader features due to stronger dipolar interactions. DP spectrum represents (blue) a combined view and quantification reference point.

The physical state of the sample further influences spectral quality. Hydrated preparations preserve native molecular organization and yield narrower linewidths, enabling better resolution of chemically distinct environments. In contrast, drying increases signal intensity due to higher sample density but results in broader resonances and reduced spectral dispersion. As a benefit the dried sample is a more quantitative means of determining chemical abundances despite broader resonances, since most signals appear in the 13C CP spectrum of this sample. These highlight the balance between sensitivity and resolution and importance of maintaining native-like conditions for accurate structural and dynamic characterization.

Quantitative comparison of CP and INEPT signal intensities further revealed differences in the relative abundance of rigid and flexible components, **Fig. 2A-D**. The ratio of CP to INEPT signal integrals was approximately ∼3-5-fold, indicating that the majority of the native wet sample is comprising flexible signals. For the isotope-labeled samples, the CP spectra of the dry sample has ∼10-fold larger S/N compared to the wet sample, due to packing efficiency differences. The wet sample is dominated by flexible chemical species, whereas the dried sample predominantly composed of rigid signal except a small amount of flexible lipid signals. We performed quantification of 13C CP signals by integration for both wet and dry sample, **Fig. 2C,D**. The results represent abundances of different fractions and chemical groups and indicate that their ratio is different for the wet and dry preparations. The 13C CP spectrum on the dry sample is more quantitative due to the conversion of most of the flexible signals into rigid ones due to drying. We utilize this for the determination, which indicates integrals of 3% for 190-160 ppm, 1% for 160-125 ppm, 69% for 125-60 ppm and 27% for 60-10 ppm of the whole signal integral, **Fig. 2C**. Compared to the quantification based on the CP spectrum on wet sample (3% for 190-160 ppm, 1% for 160-125 ppm, 48% for 125-60 ppm and 48% for 60-10 ppm,), **Fig. 2D**, the dry sample represent larger abundance of AGP, AG and PS. This indicates that the AGP fraction composed of rigid as well as flexible fractions in *M. abscessus* and requires sample drying to be quantified accurately. These abundance percentages were corrected by considering the 13C isotope-labeling efficiency that varies for different chemical groups, given as chart plots on the right side in **Fig. 2C,D**.

Finally, we recorded 1D 13C NCa/NCO spectra and compared to 13C CP to support the presence of peptide resonances and 13C signals in close proximity to nitrogen in the AGP fraction of the *M. abscessus* whole-cell sample, **Fig. 3**. These offer spectral editing and the possibility to identify the regions of peptide signals in the overlapping 1D 13C CP spectra. The 15N signal filtration, result in the peptide Ca and CO regions at ∼55 and 175 ppm. This information will be utilized along with the information from the 2D spectra to identify the chemical shifts and abundances of otherwise overlapping signals in 1D spectra.

**Figure 3.**
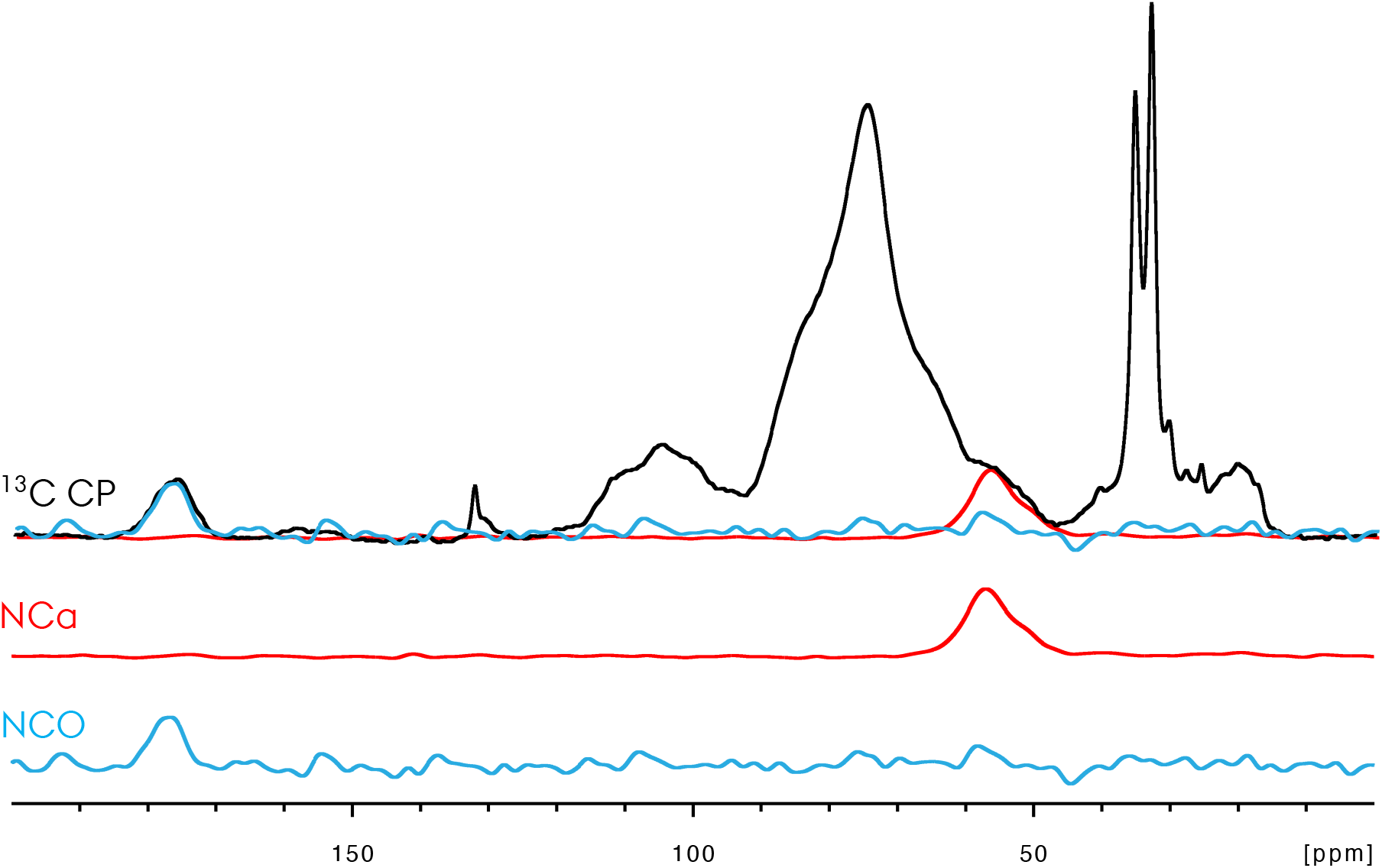
Comparison of ^13^C CP (black) ssNMR spectra with 1D ^13^C NCa (red) and NCO (blue) spectra. ssNMR spectra recorded on dry ^13^C,^15^N isotope-labeled *M. abscessus* whole-cell samples at 750 MHz recorded with 3.2 mm thin-wall rotor and 10 kHz MAS.

### 3.3. Multidimensional ssNMR at conventional 750 MHz reveals structure, dynamics and composition of *M. abscessus*

The 2D 1H-13C ssNMR correlation ssNMR experiments provide a significant increase in spectral resolution compared to 1D ssNMR methods and enable detailed analysis of chemical composition as we showed for a variety of biological planktonic and biofilm bacteria systems,^35, 41, 50, 51^ **Fig. 4A-C**. The 13C-detected 2D 1H-13C spectra were recorded via INEPT vs. CP at 10 kHz MAS, 275 K set temperature and 750 MHz. The INEPT and CP based spectra selectively probes flexible versus rigid fraction of the signals, with significantly different resonance linewidths. Both the native wet and dried samples were measured under these conditions. For the CP-based 2D 1H-13C ssNMR two different contact times (short: 0.2 ms vs. long: 4 ms) were utilized to probe different chemical connectivities within the samples and quantify morphologic signatures.

**Figure 4.**
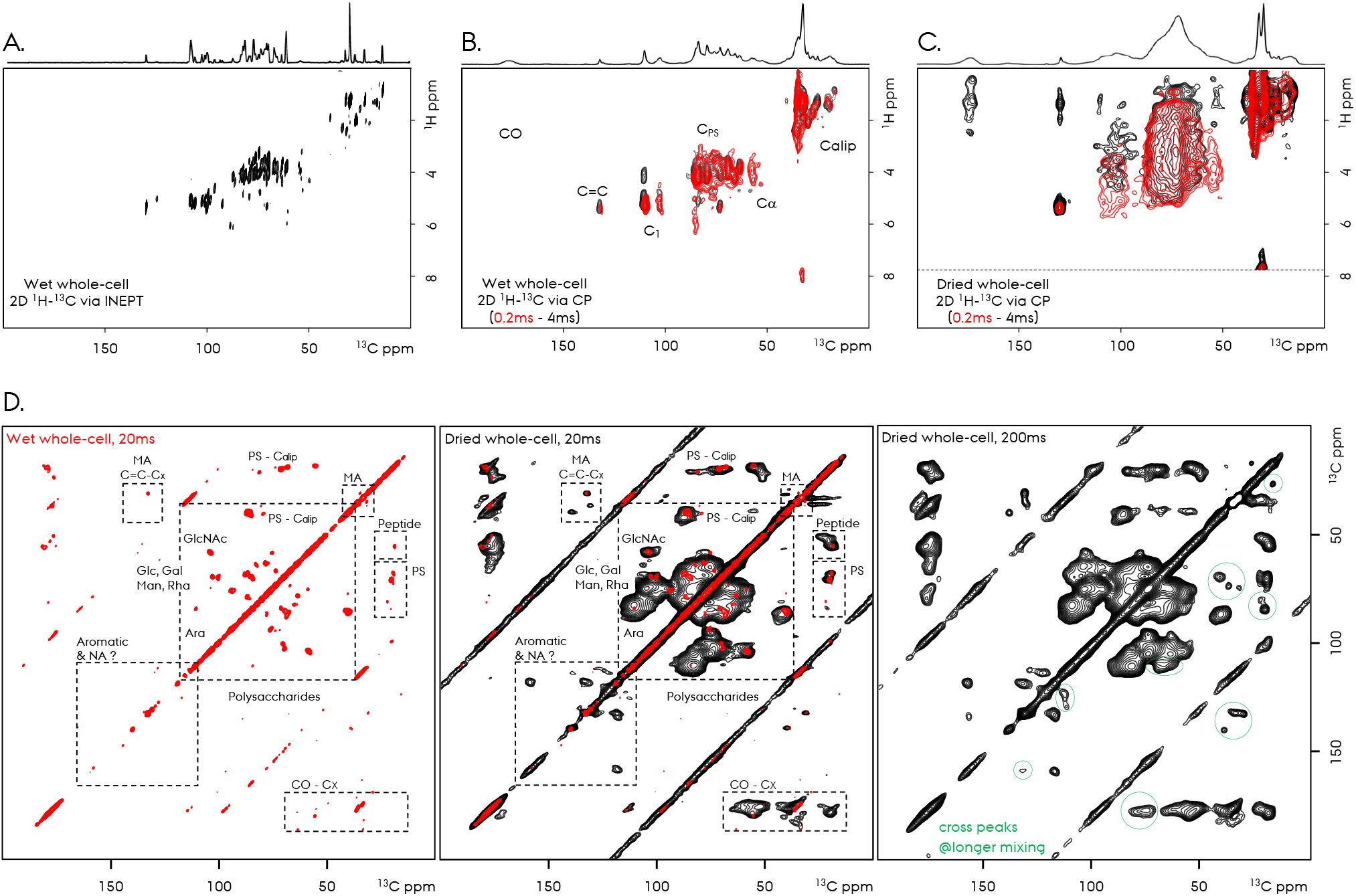
Comparison of ^13^C-detected ssNMR spectra of ^13^C,^15^N isotope-labeled *M. abscessus* whole-cell samples at 750MHz recorded with 3.2mm thin-wall rotor and 10 kHz MAS. The 2D ^1^H-^13^C spectra are recorded with INEPT/CP and ^1^H/^13^C detection. The native wet and dried *M. abscessus* whole-cell samples were recorded. See experimental section for more details. **A.** INEPT-based 2D ^1^H-^13^C ssNMR correlation spectra of wet sample. **B**. CP-based 2D ^1^H-^13^C ssNMR correlation spectra of wet sample with 0.2 and 4 ms of CP contact time. **C**. CP-based 2D ^1^H-^13^C ssNMR correlation spectra of dry sample with 0.2 and 4 ms of CP contact time. **D**. CP-based 2D ^13^C-^13^C ssNMR correlation spectra of wet/dry samples with 2 ms of CP contact time and 20/200 ms of DARR mixing.

2D 1H-13C offer well-resolved cross peaks for the wet whole-cell sample that facilitate identification of distinct molecular environments, due to the detection of the flexible signals in the INEPT based spectra, **Fig. 4A**. INEPT-based experiments selectively detect mobile components, yielding narrow linewidths and enabling assignment of signals with high confidence as we previously presented.^35, 41^ In contrast, CP-based 2D 1H-13C experiments on the wet sample detects rigid components and contain broader resonances under conventional MAS conditions, with sufficient resolution despite lower than the INEPT-based spectrum, **Fig. 4B**. Nevertheless, the CP-based 2D spectra of the wet sample comprised of well-resolved cross-peaks, **Fig. 4B**, in contract to the much broader signals obtained for the dried sample, **Fig. 4C**. The combined use of INEPT and CP provides complementary information, allowing separation of mobile and rigid fractions and revealing the distribution of dynamics across different molecular classes. In terms of sensitivity and detection of certain chemical groups, e.g. with lower abundance, remarkably, the dried sample offers greater sensitivity, despite its lower resolution, **Fig. 3C**. Many cross-peaks were observed only in these spectra and would be beneficial to consider for future ssNMR studies to characterize similar systems along with the measurements on et hydrated counterparts. 3D ssNMR spectroscopy, with 1H-detection as shown below, may resolve the broader resonances further and facilitate assignment for dry samples.

To increase the resolution and facilitate group-resonance assignment, the 2D 13C-13C correlation experiments with various DARR mixing times were recorded that provide further structural insights by establishing connectivities between 13C chemical sites, **Fig. 4D**. These experiments support our previous 2D 1H-13C based resonance assignments and help to identify specific carbohydrate motifs.^55^ More importantly, they provide information on the organization of PS networks within the cell envelope and inform on the morphology of different chemical species at the molecular level. We provide tentative group assignments in the 2D spectra as shown with dashed rectangles, **Fig. 4D**. With minimal spectral overlap compared to 2D 1H-13C, these offer a quantitative determination of chemical species, by being mindful about the different isotope-labeling efficiencies. Together, the 2D 13C-13C ssNMR spectra facilitate assignment of key spectral regions and provide a comprehensive view of chemical composition.

The 2D 1H-13C spectra with different CP contact times as well as the 2D 13C-13C ssNMR spectra with longer DARR mixing time of 200 ms, provide details of longer range chemical contacts. These spectra contain new cross-peaks, such as the ones circled in the 2D 13C-13C spectrum, **Fig. 4D**, e.g. indicating the longer range PS to CO cross peaks that are not present in the 20 ms 2D 13C-13C DARR spectra. For example, the CO-PS cross peak at ∼76/177 ppm appear only at the longer mixing time of 200 ms, that is due to the functionalized carbonyl sites in the PS chemical groups.

The 2D 13C-13C ssNMR data of wet vs. dry samples reveal differences in molecular dynamics of chemical species. Comparing the cross-peak intensities in these spectra, it’s evident that the lipid-derived signals and certain polysaccharide segments exhibit higher flexibility and dominating the 2D spectra of the wet sample, while the more rigid structural polysaccharides and peptide-associated components appear in the 2D spectra of the dried sample, **Fig. 4D**. These observations highlight the coexistence of chemical groups with differential flexibility within the NTM whole-cell sample and underscore the importance of combining different experimental approaches (dried vs. wet and CP vs. INEPT) to capture this complexity.

### 3.4 Spectral resolution and sensitivity at 750 MHz and 1.2 GHz

From the results described above, it’s clear that the dry samples (low resolution - high sensitivity) is essential to analyze, in addition to the wet samples (high resolution - low sensitivity). Some of the cross-peaks are only obtained for the dry samples when measured at 750 MHz at a 3.2 mm thin-wall rotors and 10 kHz MAS. To quantify the effect of resolution and sensitivity gain at higher magnetic fields for both wet and dry NTM samples, we recorded ssNMR spectra by 1H-detection at 1.2 GHz ultrahigh magnetic field combined with ultrafast MAS at 100 kHz utilizing 0.7 mm rotors and compared these to the 1H/13C-detcted spectra recorded at 750 MHz at a 3.2 mm thin-wall rotors and 10 kHz MAS, **Fig. 5A-C**. The influence of magnetic field strength, MAS frequency (so the ssNMR probe) and detected nucleus on ssNMR performance was evaluated. The spectra recorded at 750 MHz was recorded at 3.2 mm thin-wall rotors (∼40-50 mg sample) and 10 kHz by 1H/13C-detected, **Fig. 5A,B**. The spectra recorded at 1.2 GHz was recorded at 0.7 mm rotors (∼0.5 mg sample) and 100 kHz, **Fig. 5C**. The spectra were compared in terms of the overall sensitivity (signal/noise ratio, S/N), sensitivity per square-root of unit-time (SNT) and resolution.

**Figure 5.**
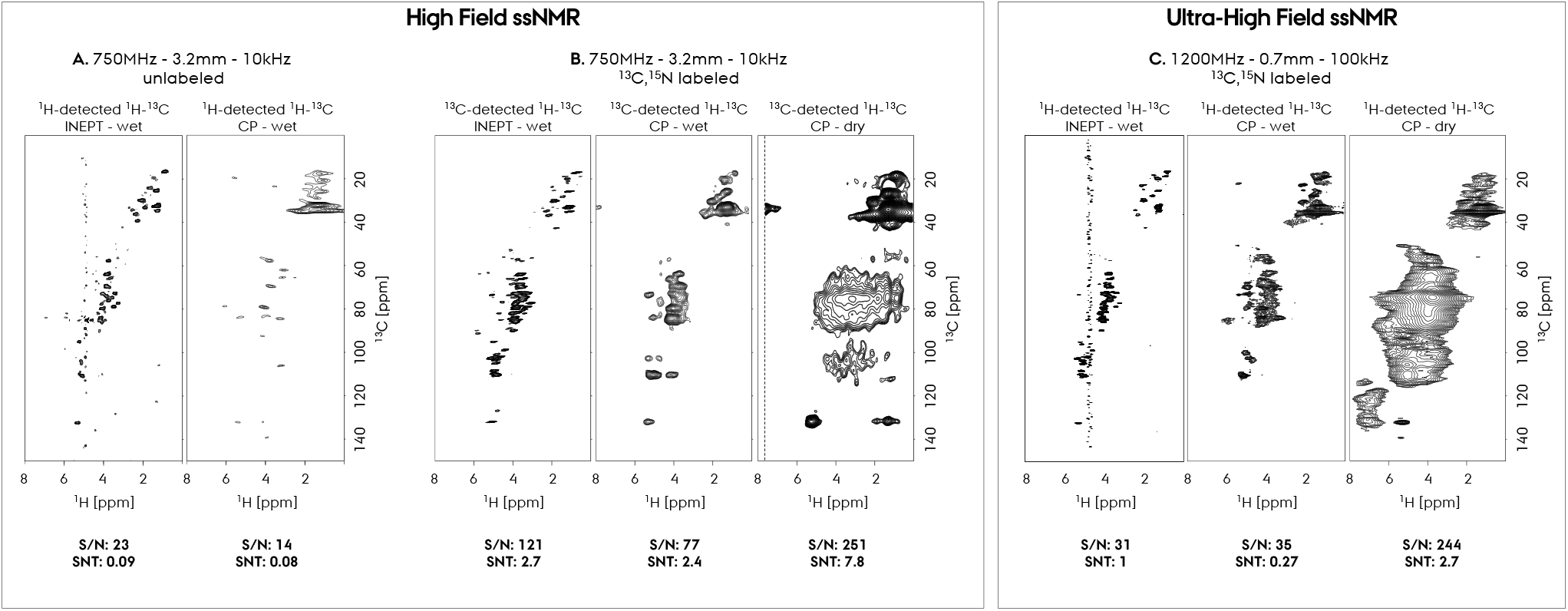
Comparison of ssNMR spectra of *M. abscessus* whole-cell samples at 750 MHz, 3.2 mm thin-wall rotor and 10 kHz MAS and 1.2 GHz, 0.7 mm rotor and 100 kHz MAS. The 2D ^1^H-^13^C spectra are recorded with INEPT/CP and ^1^H/^13^C detection. The native wet and dried *M. abscessus* whole-cell samples were recorded at natural abundance or with ^13^C,^15^N isotope-labelling. See experimental section for more details. **A)** ^1^H-detected 2D ^1^H-^13^C ssNMR correlation spectra recorded at 750 MHz, 3.2 mm probe and 10 kHz MAS with unlabeled wet sample. The 2D spectra were recorded with proton-detected INEPT and CP-based ssNMR. **B.** ^13^C-detected 2D ^1^H-^13^C ssNMR correlation spectra recorded at 750 MHz, 3.2 mm probe and 10 kHz MAS with 13C,15N isotope-labeled wet and dry sample. The 2D spectra were recorded with carbon-detected INEPT and CP-based ssNMR. **C**. ^1^H-detected 2D ^1^H-^13^C ssNMR correlation spectra recorded at 1.2 GHz, 0.7 mm probe and 100 kHz MAS with ^13^C,^15^N isotope-labeled wet and dry sample. The 2D spectra were recorded with proton-detected INEPT and CP-based ssNMR. The S/N values are given below each spectra. The S/N per unit time (SNT) values were calculated by dividing the S/N with the square-root of the total experiment time.

In our previous work, we applied 1H/13C-detected 2D 1H-13C ssNMR experiments to natural-abundance intact NTM at 750 MHz using a 3.2 mm thin-wall rotors under 10 kHz MAS.^41^ The dry samples exhibited very broader signals and reduced resolution, preventing assignment, not shown here. The same ssNMR experiments were performed on hydrated whole-cell samples with 1H-detection, where the combination of sufficient sensitivity and spectral resolution allowed tentative chemical assignments, **Fig. 5A**. The INEPT-based spectra provided well-resolved resonances from flexible regions, with a S/N:23 and SNT:0.09. However, the CP-based spectra were dominated by broad features from rigid components limiting detailed interpretation, mostly from aliphatic signals, with a S/N: 14 and SNT: 0.08. While 2D ssNMR substantially improved chemical-site identification compared to 1D, it highlights the limitations of conventional natural abundance samples and the need for higher magnetic field strengths, isotope-labeling and ultrafast MAS to resolve overlapping signal with low S/N.

To improve sensitivity, we utilized isotope-labeling and recorded similar 13C-detected 1H-13C ssNMR spectra at 750 MHz and 10 kHz MAS, as reference to be compared to 1.2 GHz experiments, **Fig. 5B**. As expected, the INEPT and CP based 2D on wet and dry isotope-labeled samples represent much better sensitivity. The INEPT-based 13C-detected 1H-13C ssNMR spectrum represent high resolution due to the detection of flexible signals, with a S/N: 121 and SNT: 2.7, compared to the CP-based spectrum, with a S/N: 77 and SNT: 2.4. Among the three spectra recorded under these conditions, the dry sample have the largest sensitivity, with a S/N: 251 and SNT: 7.8, again due to improved sample packing albeit broader resonances. As a note here, the 1H-detected spectra requires larger increments for indirect dimension, compared to the 13C-detected spectra, which compromises overall sensitivity (SNT) compared to 13-detection.

To overcome the limitations of spectral overlap and limited dispersion observed at conventional field strengths, we applied ultra-high-field ssNMR at 1.2 GHz in combination with ultrafast MAS at 100 kHz. This combination leads to a substantial enhancement in both resolution and sensitivity, representing a significant advance for the analysis of intact NTM. At 1.2 GHz, the increased chemical shift dispersion reduces overlap between signals from different molecular classes. When combined with ultrafast MAS, which more efficiently averages dipolar interactions, 1H-detected experiments exhibit markedly narrower linewidths. As a result, spectral features that appear as broad, unresolved signals at lower fields become distinguishable, allowing improved identification of lipids, polysaccharides, and protein-associated components. At 1.2 GHz, an overall improvement in spectral resolution and sensitivity was observed, manifested as narrower linewidths and enhanced separation of overlapping resonances. 1H-detection combined with ultrafast MAS at 0.7 mm rotor provide gain in sensitivity per mg sample, which is required to compensate for the sample volume reduction at 0.7 mm rotors. The amount of sample packed into a 0.7mm vs. 3.2. thin-walled rotor is nearly two orders of magnitude different, a disadvantage for the smaller rotor dimensions. Nevertheless, improved linewidth, as well as the higher magnetic field, compensated for the sensitivity loss due to much smaller sample amount. Narrower lines observed at 0.7 mm rotors result in more efficient 1H-detected spectra with larger SNT. With all these factors taken into account, the sufficient sensitivity obtained with the wet/dry NTM samples at 1.2 GHz is very encouraging. The 1H-detected 2D 1H-13C spectrum recorded at 1.2 GHz with 0.7 mm rotor for wet and dry samples result in high sensitivity, with a S/N: 31 and SNT:1 for INEPT on wet sample, S/N: 35 and SNT: 0.27 for CP on wet sample, and S/N: 244 and SNT: 2.7 for CP on dry sample. Compared to the 13C-detected 1H-13C experiments recorded at 750 MHz with 3.2 mm thin-walled rotors, the sensitivity obtained at 1.2 GHz is high, in particular for the CP experiment on dry sample which benefit from the ultra-fast MAS line narrowing effect the most.

Overall, despite the smaller overall sensitivity (SNT) values obtained at 1.2 GHz with the 0.7 mm probe, the ∼100-fold reduction in utilized sample amount, the better resolution and sufficient sensitivity is extremely valuable for future ssNMR investigations on native bacterial systems. For the difficult to observe resonances, e.g. PS signals at ∼4-6/60-120 ppm or possible nucleic acid signals at ∼7/120-140 ppm,^52^ the 1.2 GHz with 0.7 mm probe systems indispensable, since these resonances were not observed at lower magnetic field and MAS frequencies. The high resolution and sensitivity obtained at 1.2 GHz enabled acquisition of nD experiments within feasible experimental times, even for the current complex whole-cell NTM samples. Signals that were previously weak or obscured at 750 MHz became detectable at 1.2 GHz, allowing more comprehensive characterization of complex system.

Comparisons between hydrated and dried samples further highlighted the interplay between sensitivity and resolution. While dried samples produced stronger signals due to higher sample density, linewidths were consistently broader, limiting the ability to resolve closely spaced resonances at 750 MHz and 3.2 mm probe. In contrast, hydrated samples at higher field retained narrower linewidths and improved spectral clarity, demonstrating that increased field strength can compensate for reduced signal intensity in native-like conditions.

## Conclusion

In this work, we presented an ultrahigh-field nD ssNMR study on intact whole-cell *M. abscessus*. Conventional 750 MHz and ultrahigh-field 1.2 GHz MAS ssNMR experiments were combined with ultrafast MAS at 100 kHz to investigate the chemical composition, dynamics, and structural organization of *M. abscessus* under near-native conditions. Hydrated and dried whole-cell preparations were studied using isotope-labeled samples, enabling efficient nD ssNMR experiments that are otherwise not feasible at natural abundance without hyperpolarization approaches.

We utilized 1D 13C and nD 1H-13C and 13C-13C ssNMR spectroscopy to characterize the rigid and flexible fractions of the NTM cell envelope. INEPT- and CP-based experiments selectively detected flexible versus rigid molecular components and revealed pronounced differences in linewidths, sensitivity, and molecular mobility between hydrated and dried preparations. Isotope-labeling efficiencies were quantified for different chemical groups and shown to vary substantially between polysaccharides, lipids, and peptide-associated components. The multidimensional spectra enabled identification of chemical environments associated with peptidoglycan, arabinogalactan, mycolic acids, lipids, and peptide-containing regions, while long-range correlation experiments provided additional information on molecular organization and connectivity within the cell envelope.

Comparison of conventional and ultrahigh-field experiments demonstrated that the combination of 1.2 GHz magnetic field strength, ultrafast MAS, and 1H detection substantially improves spectral resolution and sensitivity despite the significantly reduced sample volume of 0.7 mm MAS rotors. Ultrahigh-field ssNMR enabled detection of weak and previously unresolved resonances, including polysaccharide and possible nucleic-acid-associated signals, that could not be clearly observed at lower magnetic field strengths and slower MAS frequencies. Together, these results establish ultrahigh-field and ultrafast-MAS ssNMR as a powerful platform for molecular-level characterization of intact NTM and provide a foundation for future investigations of cell-envelope remodeling and antimicrobial interactions in native bacterial preparations.

## Acknowledgements

UA acknowledges support from the Department of Structural Biology, University of Pittsburgh School of Medicine (UPSOM), for access to the high-field NMR and EM facilities. Start-up funding was provided by the UPSOM and a Competitive Medical Research Fund (CMRF) grant by the UPMC Health System to UA. UA also acknowledges Dr. Kristof Grohe and Dr. Sebastian Wegner for providing the image for a 1.2 GHz Bruker NMR instrument. WHD was supported by the NIH (NIAID R01-AI170607) and the Cystic Fibrosis Foundation (BOMBER21R3). We acknowledge resources from the Campus Chemical Instrumentation Center Nuclear Magnetic Resonance Facility, The Ohio State University. This facility is supported in part by funding from OSU’s Enterprise for Research, Innovation and Knowledge. The Bruker 1.2 GHz NMR instrument was funded by NSF grant RI-1 1935913. UA acknowledges Prof. Christopher P. Jaroniec for facilitating the access to 1.2 GHz NMR instrument.

